# Irrational choices via a curvilinear representational geometry for value

**DOI:** 10.1101/2022.03.31.486635

**Authors:** Katarzyna Jurewicz, Brianna J. Sleezer, Priyanka S. Mehta, Benjamin Y. Hayden, R. Becket Ebitz

**Affiliations:** Department of Neurosciences, Faculté de médecine, Université de Montréal, Montréal, QC, Canada; Centre interdisciplinaire de recherche sur le cerveau et l’apprentissage, Université de Montréal, Montréal, QC, Canada; Department of Neuroscience, Center for Magnetic Resonance Research, and Center for Neuroengineering, University of Minnesota, Minneapolis, MN, USA; Psychology Program, Department of Human Behavior, Justice, and Diversity at University of Wisconsin, Superior; Superior, Wisconsin, USA

## Abstract

We make decisions by comparing values, but how is value represented in the brain? Many models assume, if only implicitly, that the representational geometry of value is linear. However, in part due to a historical focus on noisy single neurons, rather than neuronal populations, this hypothesis has not been rigorously tested. Here, we examined the representational geometry of value in the ventromedial prefrontal cortex (vmPFC), a part of the brain linked to economic decision-making. We found that values were encoded along a curved manifold in vmPFC. This curvilinear geometry predicted a specific pattern of irrational decision-making: that decision-makers will make worse choices when an irrelevant, decoy option is worse in value, compared to when it is better. Indeed, we observed these irrational choices in behavior. Together, these results not only suggest that the representational geometry of value is nonlinear, but that this nonlinearity could impose bounds on rational decision-making.

## Introduction

Converging evidence suggests that we make decisions by gauging and comparing the values of available options. How are values represented in the brain? Early work on value encoding found value-tuned neurons: neurons whose firing rate changed essentially monotonically in proportion to the increasing value of an offer (Platt and Glimcher 1999). Since that time, a significant linear relationship between firing rate and value has become the standard test of whether or not a neuron is tuned for value (McCoy et al. 2003; Padoa-Schioppa and Assad 2006; Padoa-Schioppa and Assad 2007; Seo and Lee 2007; Lau and Glimcher 2008; So and Stuphorn 2010; Rustichini et al. 2017; Azab and Hayden 2018). However, other continuous, linear variables are not always represented via linear neuronal tuning functions, including speed (Maunsell and Van Essen 1983; Nover et al. 2005) and contrast (Ohzawa et al. 1985; Sclar et al. 1989). Indeed, neurons are often tuned for specific values of some variable–as in the case of orientation or color–and the continuous range of the variable is represented only at the population level (Hubel and Wiesel 1959; Campbell et al. 1968; Zeki 1983; Nauhaus et al. 2012; Li et al. 2014). Further, some studies have found some neurons with nonlinear tuning for value (Monosov and Hikosaka 2012; Rustichini et al. 2017; Mehta et al. 2019) and nonlinearities at the population level might explain why some studies find it difficult to discern a clear value tuning, even in regions implicated in value-based decisions (Bouret and Richmond 2010; Watson and Platt 2012). In short, although the relationship between neuronal firing rates and value is often implicitly assumed to be linear, there is reason to suspect that this hypothesis should be reexamined.

Part of the reason the linearity hypothesis has not been rigorously tested is that we have historically estimated value tuning via fitting tuning functions to the noisy responses of single neurons. Spiking noise makes it difficult to determine if a neuron truly has a nonlinear response profile, or else just happened to fire more than average for some values during a finite sampling window. However, recent advances in neural analysis techniques now allow us to take advantage of the convergence across neurons to improve our signal to noise ratio (Ebitz and Hayden 2021). By considering how value is represented across an entire population of neurons, we can directly probe the representational geometry of value within decision-making circuits with unprecedented resolution. This means that we can finally empirically determine whether the representational geometry of value is indeed linear, in line with the common assumption, or else if it takes on some higher dimensional geometry, like a curved manifold (Sohn et al. 2019; Okazawa et al. 2021) or even a high-dimensional “tangled” manifold (DiCarlo and Cox 2007; Yoo and Hayden 2018).

What are the functional implications of any of these representational geometries of value? Or, in other words, why has the linear hypothesis been so tempting? A linear system is ideally suited for value-based calculations: in linear representation, each unit change in value produces a unit change in the neuronal activity. Thus, in principle, a linear system is suitable for generating choices that satisfy axiomatic requirements of rational decisions (Rustichini et al. 2017).

However, non-linearity (and irrationality) is ubiquitous in value-based decisions, as we well know from the field of behavioral economics. Many value-based decisions are not actually rational or else are only rational under the assumption that accuracy is bounded by some kind of cognitive or computational limitations (Simonson and Tversky 1992; Lieder and Griffiths 2020). This latter idea, known as “bounded rationality” (Simon 1997; Gigerenzer 2020), is generally linked to constraints on specific cognitive capacities such as working memory or attention, but it is just as plausible that new bounds might come to light if we understood the constraints on how the brain can represent value.

Here, we first characterized the representational geometry of value then examined its consequences for rational decision-making. We focused on neurons in the ventromedial prefrontal cortex (vmPFC, area 14). Among value-sensitive regions, the vmPFC is perhaps the most strongly associated with evaluation and choice processes, and in representation of value along the kind of common scale necessary for economic decisions (Rushworth et al. 2011; Kolling et al. 2012; Levy and Glimcher 2012; Monosov and Hikosaka 2012; Strait et al. 2014). We focused on vmPFC neuronal responses to the value of offers that were encountered during a menu-search task (Mehta et al. 2019). This task was ideal for probing the representational geometry of value because offers were unidimensional, lacked any ambiguity or risk, were encountered sequentially (multiple times per trial), and offers were uniformly distributed across value space. We found that relaxing the assumption that neurons must be linearly tuned for value netted a large number of value-tuned neurons, some of which had obviously nonlinear tuning functions. At the population level, the representational geometry of value lay along an ordered, but curvilinear manifold in the neural state space, consistent with emerging evidence that curvilinear manifolds may be a ubiquitous feature of population codes for linear variables. Because of its curvature, this representational geometry predicted a specific pattern of mistakes–a specific and paradoxical type of irrational choices that we also observed in menu- search task behavior. Together, these results suggest that at least some aspects of bounded rationality may derive from constraints on neural population coding.

## Results

Two rhesus macaques performed a total of 44,335 trials (subject J: 23,826 trials; subject T: 20,509 trials) of a menu-search task (**Figure 1A**). Some analyses of this data have been presented previously (Mehta et al. 2019), however all analyses presented here are new. On each trial, a menu of 4 or 7 masked “offers” was presented. The subjects could learn about each offer by fixating it for 400 ms, at which point the mask disappeared and a reward cue was revealed. Offers were illustrated as filled bars, where the filled area reflected a magnitude of juice. Offers were uniformly distributed within the range of 0 and a maximum juice value, which differed between subjects (**Figure 1B**). When the offer was revealed, the subjects chose whether to accept it or to reject it and explore the other offers. To accept an offer, the subjects continued to fixate for an additional 300 ms. To reject, they saccaded away.

**Figure 1.**
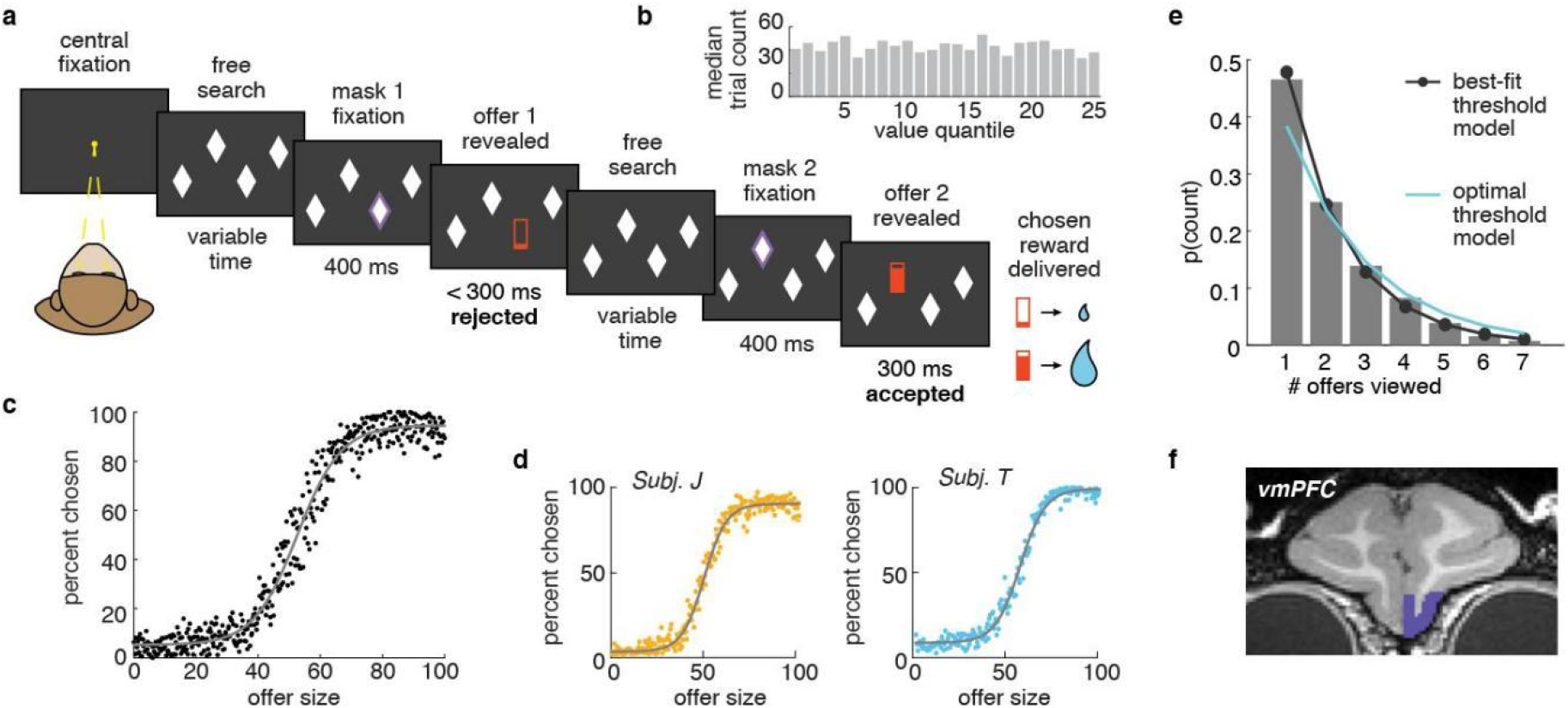
Task design and behavior. **A)** Subjects chose from menus of 4 or 7 masked offers (4 shown here). Fixating a mask (white diamond) caused it to disappear, revealing a reward cue (red bar). The filled area of the reward cue indicated juice magnitude. The subjects could freely search through the offers in any order. Subjects could accept offers by continuing to fixate them or else could reject an offer and continue to search by saccading away. **B)** Offer values were uniformly distributed between 0 and a maximum juice value. **C)** Probability of choosing an offer as a function of the value of the offer. Gray line = 4-parameter logistic curve fits. **D)** Same as **(C)**, plotted for both subjects separately. **E)** Distribution of the number of offers viewed per trial. Cyan line = prediction from an optimal compare-to-threshold model. Black line = best fit compare-to-threshold model (maximum likelihood). **F)** Recording sites in vmPFC (area 14).

Both subjects appeared to understand the task. Both chose the best offer they had seen nearly all of the time (subject J: 96% of the time, subject T: 94%) and choice behavior was well-predicted by offer values in general (**Figure 1C;** for 4 parameter logistic function [choice by value], both subjects together: slope = 14.46, intercept = 0.52, scale = 0.89, offset = 0.05, R^2^ = 0.57, n = 135250 offers; **Figure 1D;** subject J: slope = 16.71, intercept = 0.49, scale = 0.87, offset = 0.03, R^2^ = 0.58, n = 78453 offers; subject T: slope = 14.31, intercept = 0.57, scale = 0.91, offset = 0.09, R^2^ = 0.57, n = 56797 offers). Subjects also chose to evaluate something close to the optimal number of offers. On average, the subjects evaluated 2.12 offers per trial (+/− 0.25 standard deviation across trials; subject J: 2.09 +/− 0.24; subject T: 2.17 +/− 0.25; **Figure 1E**). A decision-maker implementing the optimal threshold for accepting an offer in this task (i.e. 61.6%; Mehta et al. 2019) would, on average, evaluate 2.6 offers per trial and the distribution of viewed offers would be geometric with exactly this half life (see **Methods**). Indeed, the number of offers the subjects evaluated did appear geometrically distributed, suggesting a compare-to-threshold process, though it was shifted down compared to optimal, which could indicate some impulsivity (**Figure 1E**).

### Neuronal activity in vmPFC scales with value

The responses of 122 neurons were recorded from vmPFC (**Figure 1F**; area 14; n = 70 in subject J, n = 52 in subject T). To analyze value responses in vmPFC, the trials were broken down into a series of “offer viewing periods”: the 500 ms epochs starting 100 ms after the reveal of each offer, to account for sensory processing delays (Strait et al. 2014). This epoch was chosen *a priori* to match the epoch in which the largest number of neurons were modulated by offer value in previous analyses of this dataset (Mehta et al. 2019).

The firing rates of 46/122 (38%) vmPFC neurons were correlated with offer value (**Figure 2A-D**; significantly higher mutual information between the firing rate and value, compared to shuffled value labels; significantly greater proportion than chance, p < 0.0001, one-sided binomial test). On average, across all the neurons, we found that increasing value predicted an essentially monotonic increase in the mean neuronal firing rate (**Figure 2E**; significant main effect of value bin on the firing rate, p < 0.0001, beta = 0.12, R^2^ = 0.03, linear regression, n = 3050 [25 value bins across 122 neurons]). However, the specific value-tuning functions of individual neurons were more heterogeneous than this averaged neuronal profile. Many had some curvature (**Figure 2A-B**; 19/49 [35%] of the significant neurons had curvilinear turning as indexed by being better fit by a quadratic function than a linear function, Mandel’s fitting test [see **Methods**], while 27/46 [65%] were not).

**Figure 2.**
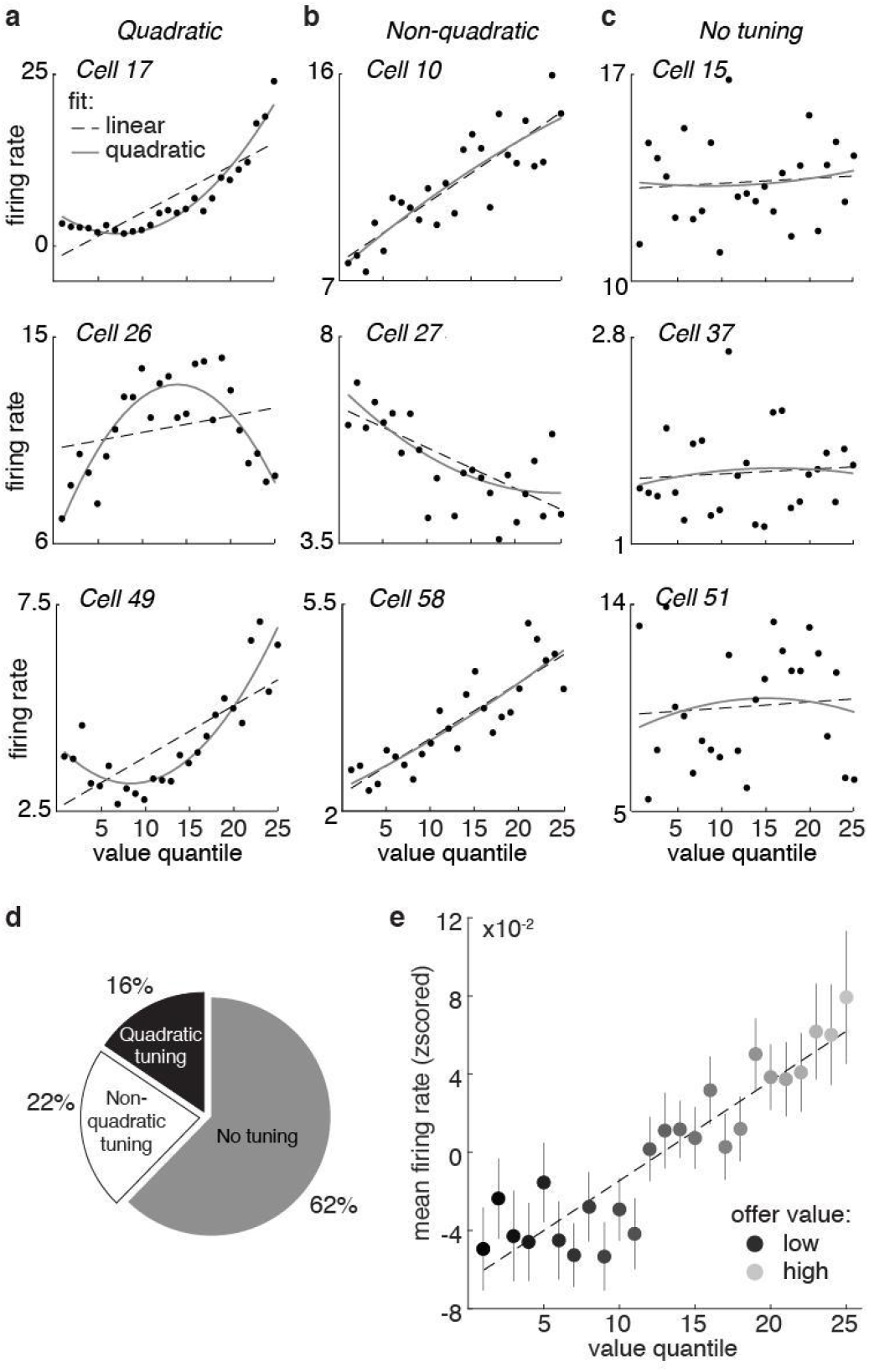
Tuning for value in individual vmPFC neurons. **A)** The firing rates of three example neurons (rows) that were quadratically tuned for value, plotted as a function of value quantile bins. **B)** Same as (A) for three example non-quadratically tuned neurons. **C)** Same as (A) for three example neurons that were not tuned for value. **D)** Proportion of all cells (n = 122) within each category. **E)** Average firing rates from all 122 neurons, plotted as a function of value quantile. Error bars indicate +-standard error of the mean across neurons (SEM).

Because we were largely interested in understanding the geometry of value coding across vmPFC neurons, we turned to population-level analyses (**Figure 3A-D**; Ebitz and Hayden 2021). The majority of the neurons were recorded asynchronously here, so the neurons were combined into pseudopopulations to perform population analyses (see **Methods**; (Meyers et al. 2008; Machens et al. 2010; Churchland et al. 2012; Mante et al. 2013; Ebitz et al. 2020). At the population level, we found that there were systematic, structured relationships between the neural responses to different offers. Offers with similar values were represented by similar patterns of neural activity–patterns that were closer together in the neuronal state space– compared to offers with different values (**Figure 3E**; significant main effect of the difference between two offers on the representational distance between the offers: p < 0.0001, beta = 2.32, R^2^ = 0.68, linear regression, n = 300 [all unique pairs of 25 value bins]). Together, these results indicate that offer values were represented in vmPFC in this dataset, both at the level of single neurons and at the level of the population.

**Figure 3.**
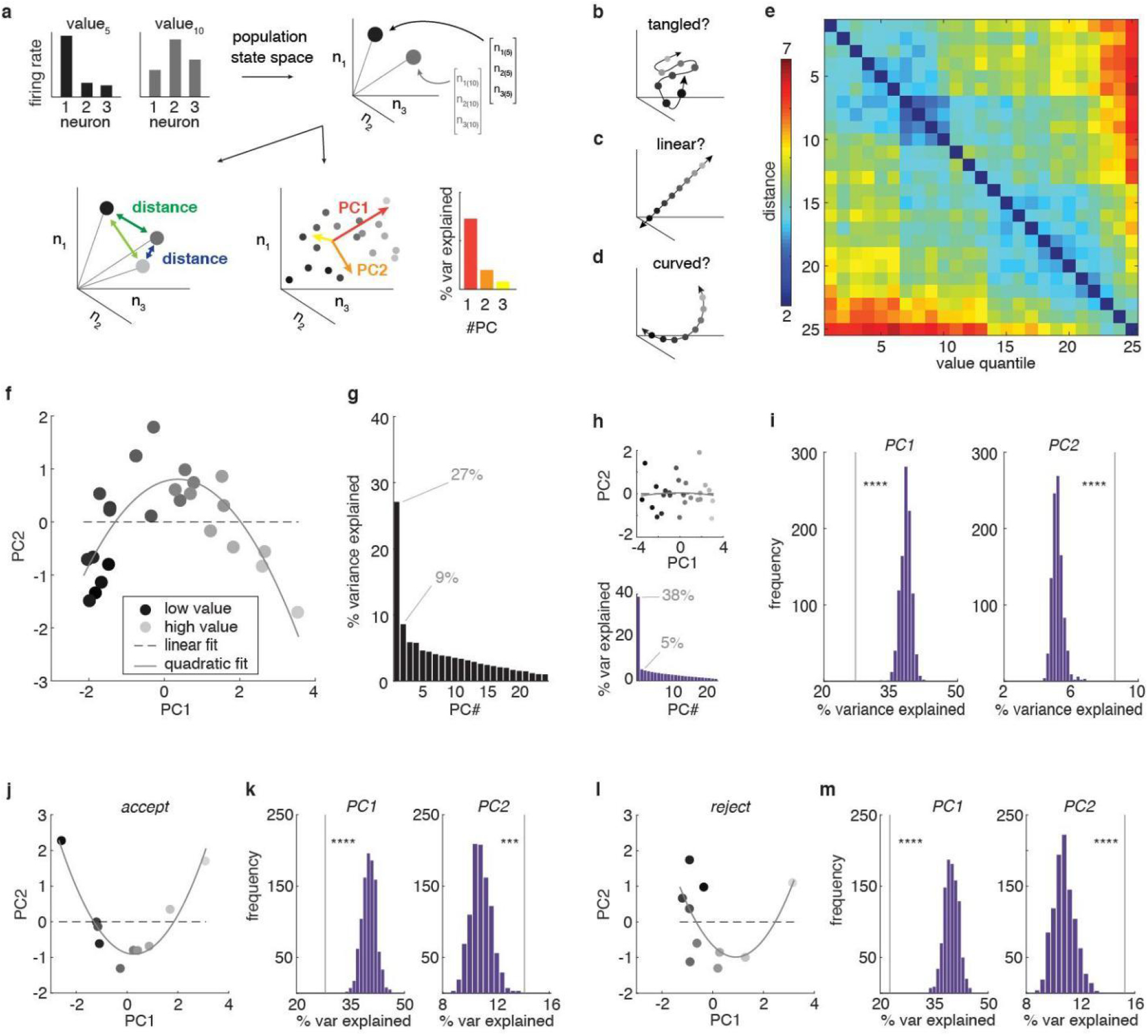
Representational geometry of value is curvilinear at the population level. **A)** The neuronal population response to an offer value is a pattern of firing rates across neurons (top left). These patterns can also be understood as vectors in a neuron-dimensional space, or, equivalently, as points in the neuronal state space (top right). We can then probe the representational geometry of these neuronal population patterns through measuring the distance between neural responses (bottom left) or via examining the major axes of covariability between neurons with principal components analysis (PCA; bottom right). **B-D)** We considered three hypotheses about how value could be represented at the population level. First, value representations could be “tangled” (B): if population value representations are very high dimensional at the population level, nearby values might not even be represented by nearby patterns of activity. Second, value representations could be “linear” (C): population value representations could follow a single straight line through the neural state space or, equivalently, occupy a single dimension in the neural state space. Third, value representations could be “curved” (D): value representations could be structured–with nearby values represented by nearby patterns of activity–but the population manifold could still occupy more than one dimension. **E)** The mean distance between neuronal states corresponding to different values. **F)** The projection of the neural population onto the first 2 principal components (PCs). Shades of gray = value bins from low (light gray) to high (dark gray). Dotted line = best linear fit. Solid line = best quadratic fit. **G)** Percent variance explained by each PC. Capturing the variance in a curved function would require more than one PC. **H)** Same as (F) and (G) for one example linearized dataset (see **Methods**). **I)** A comparison of the variance explained by the first 2 PCs in the real population (vertical line) against bootstrapped distributions of linearized datasets. Note that third and higher order PCs also continue to explain more variance in the real data compared to linearized controls (see **Results**). **J,K)** Same as **(F)** and **(I)**, but for accepted offers only. **L,M)** Same as **(F)** and **(I)**, but for rejected offers only.

### Population activity scales with value along a curvilinear manifold

The vmPFC population represented offer values in a structured way, suggesting that offer value representations were arranged in some logical order, rather than being “tangled” in some highdimensional representation in the vmPFC population (**Figure 3B**; DiCarlo and Cox 2007; Yoo and Hayden 2018). However, there were still at least 2 representational geometries that could produce this structure. For one, offer values could be arranged in a straight line, as a simple, linear sequence of neural states (**Figure 3C**). A linear geometry is thought to be important for accurate decoding: for ensuring that downstream structures can correctly and consistently infer which option is the best, no matter the precise set of options the animal is choosing between (Rustichini et al. 2017). Alternatively, offer values could fall along a curved manifold, rather than a straight line (**Figure 3D**). Recent studies find that at least some forms of perceptual information may be encoded along curved population manifolds (Okazawa et al. 2021), though it is not clear whether reward value might be encoded with this kind of geometry.

One way to arbitrate between the linear hypothesis and the curved hypothesis is to use principal components analysis (PCA). PCA is a method for reducing the dimensionality of highdimensional datasets, like neural data. It finds an orthogonal set of linear axes that explain decreasing amounts of variance in the data known as principal components (PCs). By projecting neural data onto the first few PCs, we can begin to generate a low-dimensional intuition for the structure of the high-dimensional population response. Here, projecting vmPFC population activity onto its first 2 PCs revealed a curvilinear function (**Figure 3F-G**). The shape of offer values in the reduced-dimensional-space was better described by a curved, quadratic function than a linear function (linear function: AIC = 74.21, AICc = 74.38, BIC = 77.86, n = 25, k = 3; quadratic function: AIC = 49.16, AICc = 49.71, BIC = 54.04, n = 25, k = 4; all AIC, AICc and BIC weights for the linear function <0.0001). This curvature was not apparent when offer representations were first linearized, indicating that this was not some artifact of data processing (see **Methods**; **Figure 3H**; linear function: AIC = 61.38, AICc = 61.56, BIC = 65.04, n = 25, k = 3; quadratic function: AIC = 63.43, AICc = 63.97, BIC = 68.30, n = 25, k = 4; AIC and AICc weights for the quadratic function > 0.3, BIC = 0.2; Okazawa et al. 2021).

PCA can also be used to look at curvature in the non-reduced neuronal state space: via asking how succinctly the population response can be approximated by a set of linear axes. If offer values were represented linearly, then we should be able to capture nearly all of the variance between offers with a single PC. However, in order to explain the variance in a curved function in 2 dimensions, we would need at least 2 PCs: one to describe the axis spanning the arms of the curved function and one to describe the axis of curvature. (Note that curved functions can occupy more than 2 dimensions in which case they will have more than one axis of curvature.) Therefore, determine if the data was more curved than we would expect from noise, we compared the number of PCs needed to explain the variance in the real data against linearized control populations. We found that the first PC explained significantly less variance in the real data (**Figure I**; 27.08% vs 37.66% in linearized populations, CI = [36.36%, 41.24%], p < 0.0001, bootstrapped estimate, see **Methods**), while higher order PCs explained significantly more (PC2 = 8.60% vs 5.43% in linearized populations, CI = [4.73%, 5.92%], p < 0.0001; PCs 3-4, p < 0.0001; PCs 5-14, p < 0.01; PCs 15-24, ns.). This was not an artifact of the fact that the task design required subjects to either accept or reject an offer as we observed the same pattern within both accept (**Figure J-K**, first PC: 27.97% vs 41.29% in linearized populations, CI = [37.0%, 44.78%], p < 0.0001; second PC: 14.04% vs 10.63% in linearized populations, CI = [9.61%, 12.82%], p = 0.001; PCs 3-9, p < 0.02) and reject (**Figure 3L-M**, first PC: 22.69% vs 36.86% in linearized populations, CI = [36.23%, 43.73%], p < 0.0001; second PC: 15.27% vs 12.43% in linearized populations, CI = [9.65%, 12.67%], p < 0.0001; PCs 3-8, p < 0.0001, PC 9 ns.) decisions separately. The fact that more than 1 PC was needed to explain the population structure again suggested that offers were represented curvilinearly.

### Warped decoding from the curvilinear manifold

If offer values are encoded along a curved manifold in vmPFC, it would affect our ability to accurately decode value. Linear decoders are ubiquitous in population analyses (Ebitz and Hayden 2021; Kriegeskorte and Wei 2021): they are generally computationally tractable, approximate most of the variance in many curved functions, and mirror the linear weighted sum over a population that a real downstream structure would perform (deCharms and Zador 2003; Hung et al. 2005). However, decoding accuracy would be warped in even the best linear approximation to a curved manifold (**Figure 4A**). If we compared, against their true value, values that had been encoded along a curve, then decoded from a line, we would see a subtle, but systematic compression at the tails. High values would appear lower than they actually are, and low values would appear higher. To determine if decoding accuracy was compromised in this way, we compared the predictions of a linear decoder with the true underlying values (**Figure 4B**; see **Methods**). Although inaccuracies in model fitting mean that one might see compression at the extreme values even if the underlying value coding axis was linear, the best decoder on the real vmPFC data was significantly less accurate than were decoders trained on linearized control populations (**Figure 4C**; larger root mean squared error [RMSE], real data = 3.49, linearized data = 2.82, CI = [2.64, 2.99], bootstrapped test, p < 0.0001).

**Figure 4.**
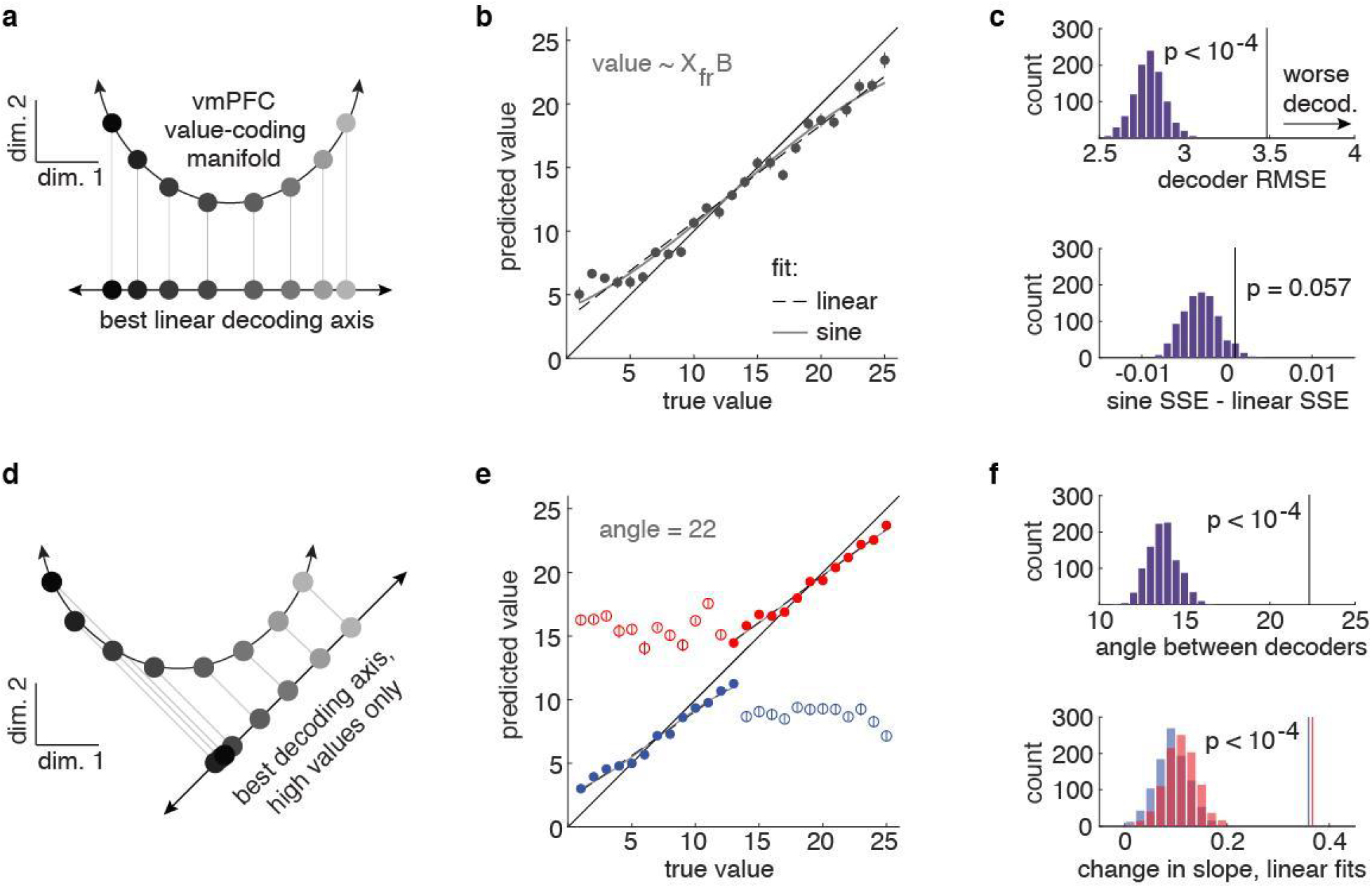
Curvilinear manifolds predict systematic biases in decoding. **A)** A curvilinear geometry would produce biases in the accuracy of a linear decoder: projection from the curve compresses the representation of values at the tails of the value range. **B)** A linear decoder trained on the population responses to values shows systematic biases in accuracy consistent with the curved shape of representation. Low values become higher, and high values become lower than the true values when predicted by the linear decoder. **C)** The best decoder on the real vmPFC data was significantly less accurate than were decoders trained on linearized control populations (top). The error in decoder accuracy may be approximated by a sine function, which describes the compression expected from encoding along the arc of a circle, visualized in (**A)**. Sine function was more likely to describe the compression observed in real data than was a linear function, when compared to the linearized control populations. **D)** Another bias predicted by a curved–but not by a linear–manifold affects out-of-range observations. A decoder trained on a portion of the curved function would not make accurate predictions about the values in the other portion of the curve. **E)** A decoder trained on the population response to one half of the values offers little information on the values outside of this range. **F)** The angle between the best linear decoder for the lower and higher halves of values was significantly larger in real vmPFC population than in linearized control populations (top), and so was the change in slope between true and predicted values (bottom).

Because the projection of an arc onto a line is a sine function, if values were encoded along a curved manifold, we might expect our residual decoding errors to follow a sine function (**Figure 4A**). In contrast, if compression is just due to inaccuracies in model fitting, residual errors should still be linear (and indeed they were in linearized populations: linear RMSE = 0.022; sine RMSE: 0.024; the linear function fit better in 897/1000 bootstrapped samples). However, in the real data, the shape of residual errors was better described by a sine function than a linear function, though this was not significantly outside the distribution of linearized populations (p = 0.057, bootstrapped test, linear RMSE = 0.037; sine RMSE = 0.036; difference between the two fits = 0.001 vs −0.003 in linearized population, CI = [-0.006, 0.002]). However, an arc might be a poor approximation to the quadratic functions that proved good approximations to the population manifold in 2 dimensions (**Figure 3F**). Although we found no closed form solution for the projection of a quadratic function onto a line, simulation suggested that we might expect the distribution of residuals in that case to asymptote vertically, rather than horizontally, consistent with what visual inspection suggested might be true here for high values at least (**Figure 4B**).

If values were represented linearly, we would be able to extrapolate a decoder trained on any portion of the manifold to predict the ordering of values on another. Conversely, in a curvilinear manifold, a decoder trained on one “arm” should offer little information at all about the values on the other arm (**Figure 4D**). Therefore, we next split the values in half and trained 2 separate linear decoders–one on the high values and one on the low values. We then used these decoders to predict the held-out low or high values, respectively. These decoders were significantly less accurate in the real data than in the linearized control populations (**Figure 4E**; high-trained, held-out low RMSE, real data = 9.94, linearized data = 7.29, 95% CI = [6.5, 8.06], p < 0.0001; low-trained, held-out high RMSE, real data = 11.50, linearized data = 6.64, 95% CI = [5.86, 7.44]; p < 0.0001). The best decoding axis for the high values and low values differed more in the real data, compared to the linearized data (**Figure 4F**; angle between decoders = 22.33; average angle between decoders in linearized data = 13.97, 95% CI = [12.27, 15.76], p < 0.0001). Further, while there was little change in the relationship between true and predicted values across test and train in linearized data (change in slope, high-trained = 0.12, 95% CI = [0.06, 0.18]; low-trained = 0.1, 95% CI = [0.04, 0.17]), this was not the case in the real data (high-trained = 0.37; low-trained = 0.36, both p < 0.0001). In fact, there was essentially no relationship between the decoder’s predicted value and the true value for out-of-range values in the real data (high-trained r = −0.01; low-trained r = −0.0002). In sum, we found systematic inaccuracies in decoding values from the vmPFC population that were consistent with the idea that values were encoded along a curved, rather than linear manifold.

### Irrational choices from a curvilinear manifold

Linear approximations to curved functions are most accurate locally, becoming perfect approximations in the limit of instantaneous segments of the curved functions. This means that when values are represented curvilinearly, the best possible linear decoder––the upper bound on how accurately a downstream neuron could decode value–would be sensitive to the range of values. For narrow ranges, decoding would be fairly accurate, and nearby values easy to discriminate (**Figure 5A**). Conversely, as the range of values increases, decoded values at the tails of the curved manifold would become compressed and easily confused. In the context of economic decision-making, this predicts a specific violation of one of the principles of rational choice theory: the independence of irrelevant alternatives axiom (Arrow 1951; von Neumann and Morgenstern 1953; Ray 1973). Rational decision-making requires that choices between good offers not be affected by the availability of low-value offers known as “decoys”: the choice between 2 good offers should not depend on the value of a third option, provided the value of the third option is worse than the value of both good options. However, by increasing the range, decoy offers would warp the upper bound on decoding accuracy, meaning that they could, in theory, compromise the subjects’ ability to accurately discriminate between high-value offers.

**Figure 5.**
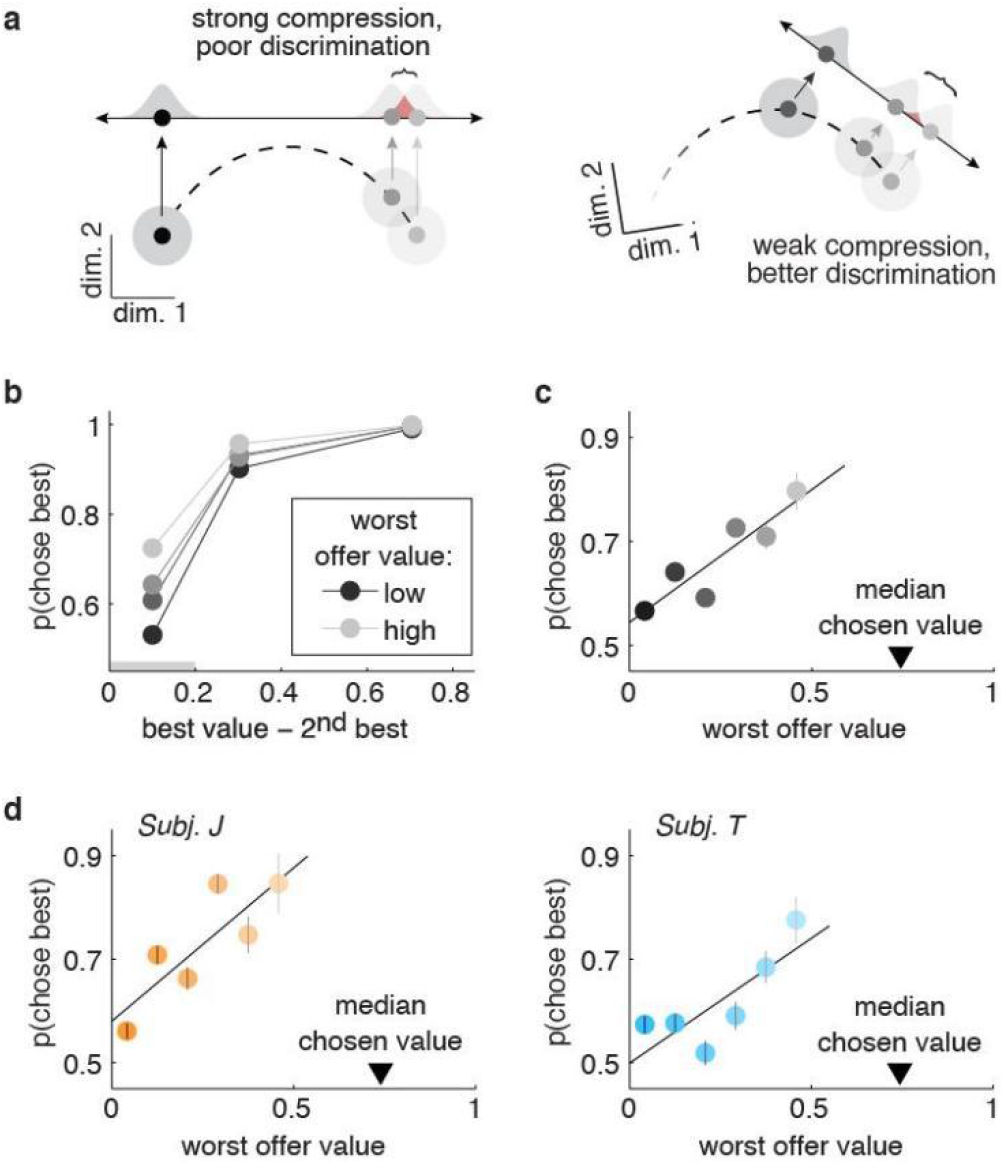
Curvilinear manifolds suggest a specific pattern of irrational decisions. **A)** A cartoon illustrating how the ability to discriminate good offers could depend on how bad the worst offer is in a curved manifold. Note that the linear decoder illustrated here represents an upper bound on how accurately a downstream neuron could decode value. **B)** Probability of choosing the best option seen within each trial as a function of a difference between the best and second best option, plotted separately for different values of the worst option (shades of gray). The gray bar on the x-axis indicates the range of best-2^nd^ best values that was chosen for subsequent analyses, based on the fact that subjects only confused good offers that were within this range. **C)** Probability of choosing the best offer seen as a function of the worst offer seen, only for trials in which the best and 2^nd^ best offers were within 20% of each other. Line = GLM fit. **D)** Same as **(C)**, plotted for both subjects separately. Error bars in each graph indicate +-standard error of the mean across sessions (SEM). These are sometimes smaller than the symbols.

In general, subjects were better at choosing the best option as it got larger in comparison to the second-best option (**Figure 5B**). However, when the values of the best and second-best options were close together, and presumably most difficult to discriminate (see **Methods**), the worst option in the set systematically altered their ability to choose the best option (**Figure 5C**). When the worst option was very low, the subjects’ ability to discriminate between the best and second- best option approached chance (sig. effect of decoy value: p < 0.0001, beta = 1.70; sig. effect of best-next, p < 0.0001, beta = 12.61). This effect was significant in both subjects individually (**Figure 5D**; subject J: decoy value, p < 0.0001, beta = 2.68; best-next, p < 0.0001, beta = 14.22; subject T: decoy value, p < 0.0002, beta = 1.33; best-next, p < 0.0001, beta = 11.10). This was not due to some paradoxical tendency to choose the decoy option more when its value was lower (n. sig effect of decoy value on p(choose decoy), p = 0.95, beta = −0.03; sig. effect of best- next, p < 0.0002, beta = −3.87). Subjects were more likely to see very low decoy option when they viewed more offers in a trial (correlation between the value of the decoy option and the set size, r = −0.33, p <0.0001) but the set size could not account for the decrease in accuracy: when we subselected only those trials in which subjects viewed three options, the effects were the same (sig. effect of decoy value, p < 0.0001, beta = 1.86; sig. effect of best-next, p < 0.0001, beta = 14.54; subject J: decoy value, p < 0.0005, beta = 2.38; best-next, p < 0.0001, beta = 13.30; subject T: decoy value, p < 0.0003, beta = 1.74; best-next, p < 0.0001, beta = 15.03). Accounting for other potential confounds such as the order or recency of the worst option could not explain the effect (parameter estimates were very similar and still highly significant in a model which also included set size, the order of the decoy option and its recency with respect to the best option: sig. effect of decoy value: p < 0.0001, beta = 1.26; sig. effect of best-next, p < 0.0001, beta = 12.99). In sum, what should have been an irrelevant decoy–the worst option in the set–interfered with decision-making the most when it was furthest from the best available options, exactly as we would predict if the values were decoded from a curved manifold.

## Discussion

Value is the key variable of economic decision making and its representation in the cortex likely has consequences for value-based decisions. In vmPFC, a region causally implicated in value- based decision-making (Noonan et al. 2010; Camille et al. 2011; Noonan et al. 2017), we found that the average firing rates across neurons scaled positively and linearly with value. However, many individual neurons were non-linearly tuned for value and, at the population level, the patterns of neuronal activity that encoded value traced a curvilinear manifold, rather than a straight line. This curvilinear geometry could explain why not every vmPFC study finds robust value tuning (Bouret and Richmond 2010; Watson and Platt 2012): random samples of neurons are low-dimensional projections of the underlying population geometry and a curvilinear value manifold would have many low-dimensional projections that would lack even monotonic tuning with respect to value (Okazawa et al. 2021). The neuronal code for value has long been assumed to be linear and at least quasi-linear tuning for value has been reported in many brain regions (McCoy et al. 2003; Padoa-Schioppa and Assad 2006; Padoa-Schioppa and Assad 2007; Seo and Lee 2007; Lau and Glimcher 2008; So and Stuphorn 2010; Rustichini et al. 2017; Azab and Hayden 2018). However, our results reinforce the idea that single neurons can be meaningfully tuned for value without being linearly tuned for value (Monosov and Hikosaka 2012; Mehta et al. 2019). They also suggest that the population-level representational geometry of value can be strongly nonlinear, even when the average response across neurons is not.

If a set of option values is represented linearly, it is trivial for a downstream region to decode these values in a way that respects fundamental axioms of rational value-based decisionmaking, like transitivity, completeness, and the independence of irrelevant alternatives. The last of these, independence of irrelevant alternatives, is the dictum that decisions between good alternatives should be unaffected by the value of low-value, decoy options (Arrow 1951; von Neumann and Morgenstern 1953; Ray 1973). If the representation of value were curved, however, decoys would affect decision-making. Because curvature both (1) warps the representation of values, and (2) scales positively with the range of values, the lowest-value decoys would systematically limit the upper bound on the discriminability of high-value alternatives. Indeed, we found that subjects exhibited exactly the pattern of choices predicted by their curvilinear manifold: the lower the value of the decoy option in the set, the less accurate the subjects were at choosing among better offers. This result resonates with a broad literature showing that violations of the independence of irrelevant alternative axioms are surprisingly common in real-world decision-makers (Doyle et al. 1999; Shafir et al. 2002; Choplin and Hummel 2005; Louie et al. 2013; Chau et al. 2014; Khaw et al. 2017; Chau et al. 2020). However, note that high-value decoys can sometimes compromise decision-making more than low-value decoys (Louie et al. 2013; Chau et al. 2020), perhaps because divisive normalization effects emerge when offers are presented simultaneously, rather than sequentially (Khaw et al. 2017). Future work is needed to fully understand the consequences of curvilinear value encoding for rational decision-making.

The links we have drawn so far between the curved value-coding manifold and irrational decision-making may seem predicated on the notion that a downstream region would decode value in a way that is essentially linear and, more curiously, flexible–changing across trials in response to the range of values. Linear decoders have a nice parallel to the weighted sum over neurons that a downstream neuron would have access to (deCharms and Zador 2003; Hung et al. 2005; Kriegeskorte and Diedrichsen 2019) and both neuronal activity (Abbott 1994; deCharms and Zador 2003; Chang and Tsao 2017) and behavior (Hong et al. 2016; Sohn et al. 2019) are often well described as a linear decoding of some input, even when that input is curvilinear (Sohn et al. 2019). Further, though this study cannot say what decoding scheme might be used by any downstream region(s), linear functions offer a good approximation for many nonlinear functions so the true decoding scheme need not be strictly linear for decoding accuracy to be systematically warped. The second, more provocative premise underlying our hypothesis is the idea that decoding schemes could be flexible. In order to be affected by the decoy, a downstream region would have to have the capacity to flexibly alter its decoding strategy in order to maximize the information available about the currently available set. Although it is not clear if the brain is capable of such flexibility, neural representations certainly drift over time (Rule et al. 2020; Montijn et al. 2016) and change according to both internal states and task contexts (Çukur et al. 2013; Ebitz and Platt 2015; Sohn et al. 2019; Ebitz et al. 2019; Sasaki et al. 2020) in such a way that flexible decoding may be essential. A flexible decoding scheme could also give a mechanistic explanation for other irrational biases in decision-making, like anchoring effects: the tendency of the first offer seen to shape how subsequent offers are evaluated. However, future work is needed to determine how the range of offer values shapes the information available to regions downstream of vmPFC.

If decoding were not flexible, encoding values with a curvilinear representational geometry might still affect decision-making. This is because in a fixed, quasi-linear decoding from a curved manifold, the separation between nearby values is maximal at the center: at precisely the values that are closest to the accept/reject threshold the animals adopted here. In fact, because the difficulty of the accept/reject discrimination is aligned with the axis of curvature, another possible interpretation of this result is that vmPFC encodes difficulty as well as value. However, we would argue that it is more parsimonious to imagine that vmPFC “encodes difficulty” as a byproduct of the inherent curvature in the representation of value. Curvature emerges naturally from fundamental constraints on the dynamic range of neuronal firing rates (i.e. the absolute lower bound at zero and an upper bound at the neuron’s metabolic limits; Okazawa et al. 2021) and appears even in tasks where there is no natural alignment with any difficulty axis (Sohn et al. 2019; Sabatini and Kaufman 2021). Further, even when the axis of curvature aligns with task difficulty, position along the “difficulty” axis does not appear to predict animals’ self-reported confidence in their decisions (Okazawa et al. 2021). However, just because information is not used for a specific metacognitive judgment does not mean that it is not used. Future work is needed to determine if the apex of the curvature is sensitive to changes in threshold or else if other neural correlates of decision difficulty could be an artifact of curved value representations (Kolling et al. 2016; Ebitz and Hayden 2016).

The idea that curvature can exist even in a circumstance where it compromises behavior lends credence to the notion that curvature is a ubiquitous feature of the population encoding of any linear variable. This study stands alongside growing evidence for curved representational geometries across several brain structures for several linear variables, including those underlying perceptual decisions (Okazawa et al. 2021), motor control (Sabatini and Kaufman 2021) and interval reproduction (Sohn et al. 2019). Given a bounded dynamic range of neuronal firing rates, curvature could be a powerful way to maximize information coding (Okazawa et al. 2021). Though linearity might be essential for rational decision-making, a linear information- coding manifold in a bounded space represents a significant compromise in the discriminability of states along its length compared to coding manifolds that have more dimensions (DiCarlo and Cox 2007; Fusi et al. 2016; Yoo and Hayden 2018; Okazawa et al. 2021). As the curvature of information-coding manifolds expands into more dimensions, the number of representations that can be discriminably encoded along that manifold will also increase (Ebitz and Hayden 2021). However, it is not yet clear whether the curvilinear value manifold reflects a functional signal or exists as a by-product of constraints on neuronal firing rates. Of course, these ideas are not mutually exclusive. In either case, the curvilinear geometry of value could represent a critical bound on rational decision-making that will help shape future models of choice behavior.

## Methods

All procedures were designed and conducted in compliance with the Public Health Service’s Guide for the Care and Use of Animals and approved by the University Committee on Animal Resources at the University of Rochester. Subjects were two male rhesus macaques (*Macaca mulatta*: subject J age 10 years; subject T age 5 years). Initial training consisted of habituating animals to laboratory conditions, to head restraint, and then to perform oculomotor tasks for liquid reward. Standard surgical techniques, described previously (Strait et al. 2014), were used to implant a small prosthesis for holding the head and Cilux recording chambers (Crist Instruments) over the vmPFC. Position was verified by magnetic resonance imaging with the aid of a Brainsight system (Rogue Research Inc.). After all procedures, animals received appropriate analgesics and antibiotics. Throughout all sessions, the chamber was kept sterile with regular washes and sealed with sterile caps. Some analyses of these data have been included in earlier work (Mehta et al. 2019). All analyses presented here are new.

### Recording sites

We approached vmPFC through a standard recording grid (Crist Instruments), guided by a Brainsight system and structural magnetic resonance images taken before the experiment. Neuroimaging was performed at the Rochester Center for Brain Imaging, on a Siemens 3T MAGNETOM Trio Tim using 0.5 mm voxels.The accuracy of the Brainsight guidance was confirmed by listening for characteristic differences between white and gray matter during electrode penetrations. Gray-white matter transitions occurred at the penetration depths predicted by the Brainsight system in all cases. We defined vmPFC according to the Paxinos atlas (Paxinos et al. 2000). This meant we recorded from a region of interest lying within the coronal planes situated between 42 and 31 mm rostral to interaural plane, the horizontal planes situated between 0 to 7 mm from the brain’s ventral surface, and the sagittal planes between 0 and 7 mm from medial wall. This region is within the boundaries of Area 14 according to the atlas. Electrophysiological techniques, eye tracking and reward delivery

A microdrive (NAN instruments) was used to lower single electrodes (Frederick Haer & Co., impedance range 0.7 to 5.5 MU) until waveforms of between one and four neurons were isolated. Individual action potentials were isolated on a Plexon system (Plexon, Inc.). We only selected neurons based on their isolation quality; never based on task-related response properties. All collected neurons for which we managed to obtain at least 300 trials were analyzed. In practice, 86% of neurons had over 500 trials (this was our recording target each day).

An infrared eye-monitoring camera system (SR Research) sampled eye position at 1,000 Hz, and a computer running Matlab (Mathworks) with Psychtoolbox and Eyelink Toolbox controlled the task presentation. Visual stimuli were colored diamonds and rectangles on a computer monitor placed 60 cm from the animal and centered on its eyes. We used a solenoid valve to control the duration of juice delivery, and established and confirmed the relationship between solenoid open time and juice volume before, during, and after recording.

### Experimental Design

Subjects performed a menu-search task (**Figure 1A**) that has been described previously together with subjects’ previous training history (Mehta et al. 2019).

To begin each trial, the animal fixated on a central dot (50 ms), after which either four or seven white diamonds (“offers”) appeared in randomly selected, non-overlapping positions on the screen. The number of offers per trial was either 4 or 7, and was chosen at random on each trial. Continuous fixation on one diamond for 400 ms caused it to disappear and reveal a reward offer. Offers were orange bars, partially filled-in to indicate the value of the riskless offered reward. The percentage of the offer bar that was filled in corresponded to the offer value in terms of percent of the maximum value possible per offer (e.g. an offer bar that was 10% orange and 90% black would indicate an offer worth 10% of the maximum value; 20 uL for subject T and 23 uL for subject J). Reward values for each offer were generated randomly from a uniform continuous distribution ranging from 0%-100% of the maximum possible reward value for the individual subject (although continuously varying rewards were generated, the need to represent offers in pixels did discretize offers into approximately ~200 steps).

The subject could freely search through the diamonds in any order and could accept any offer. Acceptance led to the end of the trial; rejection led to a return to the initial state (viewing an array of diamonds). To accept the offered reward, the animal had to maintain fixation on the offer for 300 ms, after which the screen would go black and the offered amount of liquid reward would be delivered immediately. Thus, selecting a given offer required 700 ms: 400 to unmask it and an additional 300 to obtain it. If the subject broke fixation on the reward stimulus at any point between 0 and 300 ms from the initial reveal, the reward stimulus would disappear and the diamond would return in its place (a “rejection” of the offer). The subject could then resume freely inspecting other offers. There was no limit to how many offers a subject could inspect, nor to how many times a subject could return to re-inspect a particular offer. The trial only ended (and a liquid reward was only delivered) after the subject accepted an offer. Reward delivery was followed by a 4 second inter-trial interval.

### General Data Analysis Techniques

Data were analyzed with custom software, written in Matlab. Neural activity was analyzed in the fixed 500 ms epoch beginning 100 ms after the offer was revealed. This epoch was chosen *a priori*, based on a previous publication (Mehta et al. 2019).

### Analysis of Choice Behavior

The menu search task was inspired by optimal stopping problems, like the well-known “secretary problem” (Freeman 1983). This means that the task encouraged the subjects to balance the goal of choosing the best offer possible against the time costs of evaluating each offer. One reasonable strategy in optimal stopping problems is to compare each offer against some fixed threshold, then choose the first offer that exceeds this threshold. Because offers were uniformly distributed in value and presented at random, there was an identical probability that any offer would exceed a fixed threshold, and that probability would be exactly equivalent to 1 minus the threshold. Because the distribution for a stopping process with a fixed probability of stopping at discrete step is a geometric one, the number of options evaluated by a decisionmaker pursuing a compare-to-threshold strategy would also be geometric. The maximum likelihood fit of a geometric distribution to the subjects’ behavior is illustrated in **Figure 1E** (model fit via the expectation maximization algorithm). The geometric distribution has a single free parameter, which can be expressed equivalently as 1 minus the probability of stopping (here, threshold), or its inverse: the average duration of the stopping process (here, number of offers evaluated per trial). Previously, we calculated the optimal, reward-rate maximizing threshold in this task (Mehta et al. 2019). Here, we used these mathematical insights to illustrate the distribution over the number of offer evaluations that we would expect from this optimal threshold. Although the geometric distribution was a good fit to the data and the subjects’ curve was close to optimal, this task does differ from the classic “secretary problem” because the subjects could return to, re-evaluate, and choose a previously-rejected offer at any time.

To determine how well choice behavior was predicted by offer values, we fit a four-parameter logistic model. The probability of accepting an offer was modeled as a function of the value of that offer according the following equation:

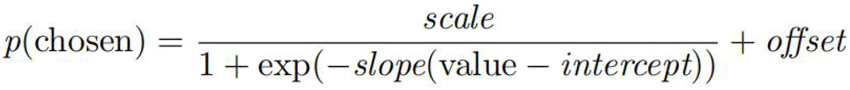

Where “value” was the objective value of the offer on the screen, the intercept captured the value where choices were evenly split between accept and reject, the slope reflected the noise around this intercept (i.e. decision temperature), and the scale and offset reflected the tendency to reject or accept offers by chance, respectively. The model was fit with the default settings of the Matlab fit.m function (minimizing nonlinear least squares, trust region algorithm) and the adjusted R^2^ was taken as an index of the quality of model fit.

### Neuronal Tuning

To identify which neurons were tuned for value, we used an approach that makes no assumptions about the shape of neuronal value tuning. Specifically, we calculated mutual information between the firing rate and the values for each individual neuron, and compared it against a distribution in which the labels had been shuffled (n = 1000 shuffles). Firing rates and values were both divided into quantile bins, where the number of bins (3 each) was chosen to minimize the number of empty cells (unobserved combinations of firing rate and value) while still allowing for tuned neurons with U-shaped value tuning functions. We wanted to avoid empty cells because these can inflate mutual information estimates: it is not clear whether the probability of an empty combination of variables is truly zero or if it simply was not observed within the finite sample of data.

To determine the shape of tuning functions within the set of tuned neurons, we used Mandel’s fitting test (Mandel 2012; Andrade and Gómez-Carracedo 2013). This method asks if a mean firing rate model that permits curvature (i.e. includes a quadratic term) is a significantly better fit than a model that does not (i.e. a model that includes only linear terms) via calculating the following F-statistic:

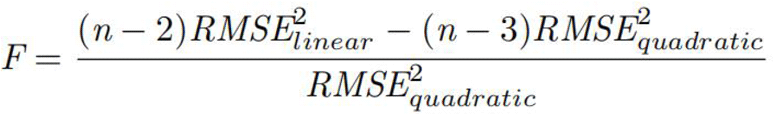

Where RMSE stands for root mean squared error of the linear and quadratic regression models, respectively, and n is the number of value bins. Because this analysis assumes normally distributed data, offers were quantile-binned into 25 distinct values and the models were fit to average firing rates in each bin.

### Pseudopopulations

Neurons in this study were recorded largely separately, so to gain insight into the representational geometry of value coding at the population-level, we built pseudopopulations from non-simultaneously recorded neurons (Meyers et al. 2008; Machens et al. 2010; Churchland et al. 2012; Mante et al. 2013; Ebitz et al. 2020). The pseudopopulation approach does not permit a reconstruction of the covariance structure between simultaneously recorded neurons, but it can still be useful for generating first order insights into how population activity changes across various conditions.

Offers were first quantile-binned into 25 distinct values, a level at which each value bin spanned a +/− 2% change in reward. Within each bin, firing rates from separately recorded neurons were randomly drawn with replacement to create a pseudotrial firing rate vector, with each entry corresponding to the activity of one neuron when an offer within that quantile bin was on the screen. Pseudotrial vectors were then stacked into a trials-by-neurons pseudopopulation matrix. Twenty-eight pseudotrials were drawn from each cell for each condition, because at least 95% of pairs of cells and conditions had at least this number of observations. One neuron was excluded because it did not spike within the selected epoch. All pseudopopulation results are reported for a single, randomly-seeded pseudopopulation, but were later confirmed with a range of different random seeds. For analyzing only accepted and only rejected offers, accept-only and reject-only offers were quantile-binned into 10 distinct values. The number of bins was chosen to account for the smaller number of observations per bin when the data was split according to choice (95% of pairs of cells and conditions contained at least 28 observations, as for the analysis of all choices together).

### Linearized Control Populations

In order to determine if the apparent curvature in vmPFC populations was just an artifact of some data processing step(s), we generated 1000 bootstrapped estimates of the distributions of certain statistical measures under the null hypothesis that the neural representation of value was linear. We did this via linearizing pseudopopulation responses (after Okazawa et al. 2021). A line was fit to the neural population data, then interpolation was used to identify an evenly spaced set of neural states along that line. To generate realistic trial-to-trial noise, spiking observations were then drawn from poisson distributions parameterized by the neural states.

### Decoding Models

To decode offer values from the neural population data, we fit general linear models of the form:

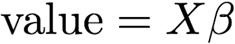

Where X is the matrix of population data (n-trials by k-neurons), augmented by a column of ones to serve as an intercept, β is a vector of weights for each neuron (or intercept), and value is one of 25 value bins, with the range rescaled between 0 and 1. Decoding models were trained via maximum likelihood using standard Matlab libraries (glmfit). Decoding was performed via projecting the real data onto the optimized β vector. In order to maximize the accuracy of the models’ predictions–and thus the accuracy of our estimate of their residual errors–decoding models were trained and tested on complete data unless otherwise specified.

### Decoy Effects

To determine whether choices were irrational in a way that matched the predictions of the curvilinear value manifold, we asked if the value of an ostensibly irrelevant decoy option altered the subjects’ choices. In this task, subjects chose between 4 or 7 options, but were free to evaluate as many (or as few) options as they wanted on each trial. To conduct this analysis, we therefore first isolated the trials in which subjects chose to view at least 3 separate options (16,933 total trials; subject J: 9,196; subject T: 7,737; 26.5% of trials), because these were the only trials in which the subject had seen a non-overlapping best option, second-best option, and a worst option. Subjects had little difficulty choosing between the best and second-best options when the difference between their values was large: subjects chose the best option 99% of the time when its reward was at least 0.2 units greater than the second-best, but only 70% of the time within this range. Therefore, we focused our analyses on the trials where the difference between the best and second-best option were within 0.2 (8,157 total trials; subject J: 4,037; subject T: 4,120). To determine whether the value of the irrelevant decoy option affected the probability of choosing the best option, we fit a GLM that included the main effects of the decoy option’s value and the difference between the best and second-best options. The probability of choosing the best option was modeled as a Bernoulli random variable. As a control for any effects of the set size, all analyses were repeated within the subset of trials in which subjects had seen only three unique options (2,806 total trials; subject J: 1400; subject T: 1196). As an additional control, we also fit a GLM that included terms for the set size (3-7), the order that the decoy option was presented (first, second, etc.), and the recency of the decoy option with respect to the final choice (one-back, two-back, etc.). These additional variables were generally significant, but did not substantially alter the parameter estimates or significance of the effects of interest here (i.e. the effect of the decoy offer value on choice accuracy).

